# Neural Representations of Faces are Tuned to Eye Movements

**DOI:** 10.1101/402263

**Authors:** Lisa Stacchi, Meike Ramon, Junpeng Lao, Roberto Caldara

## Abstract

Eye movements provide a functional signature of how human vision is achieved. Many recent studies have reported idiosyncratic visual sampling strategies during face recognition. Whether these inter-individual differences are mirrored by idiosyncratic *neural* responses has not been investigated yet. Here, we tracked observers’ eye movements during face recognition; additionally, we obtained an objective index of neural face discrimination through EEG that was recorded while subjects fixated different facial information.

Across all observers, we found that those facial features that were fixated longer during face recognition elicited stronger neural face discrimination responses. This relationship occurred independently of inter-individual differences in fixation biases. Our data show that eye movements play a functional role during face processing by providing the neural system with information that is diagnostic to a specific observer. The effective processing of face identity involves idiosyncratic, rather than universal representations.

## INTRODUCTION

The visual system continuously processes perceptual inputs to adapt to the world by selectively moving the eyes towards diagnostic information. As a consequence, eye movements do not unfold randomly, and during face processing humans deploy specific gaze strategies. Since Yarbus’s seminal report (Yarbus, 1967), a multitude of studies have reported a distinct triangular fixation pattern encompassing the eye and mouth region during face recognition (Henderson, Williams, & Falk, 2005). For many years, this T-shaped fixation pattern was considered to be universal and shared across observers, suggesting that the presence of a face triggers a unique biologically-determined information extraction pattern.

However, over the last decade, a growing body of work has challenged this view by revealing cross-cultural (e.g., Blais, Jack, Scheepers, Fiset, & Caldara, 2008; Miellet, Vizioli, He, Zhou, & Caldara, 2013), idiosyncratic (Mehoudar, Arizpe, Baker, & Yovel, 2014) and even within-observer (Miellet, Caldara, & Schyns, 2011) differences during face recognition. Contrary to the average T-shaped fixation pattern displayed by Westerners, Easterners deploy a global sampling strategy by directing the majority of their fixations toward the center of the face, while reaching comparable efficiency in face recognition (for a review see Caldara, 2017). Even among Western observers there is a conspicuous degree of variability in the sampling strategies adopted to achieve face identification (Miellet et al., 2011). In addition, in line with early observations based on individual participants (Walker-Smith, Gale, & Findlay, 1977), recent studies demonstrate that observers deploy unique sampling strategies (Kanan, Bseiso, Ray, Hsiao, & Cottrell, 2015; Arizpe, Walsh, Yovel, & Baker, 2017), which are stable over time (Mehoudar et al., 2014), and relevant to behavioral performance (Peterson & Eckstein, 2013). Individuals’ sampling strategies deviate considerably from the well-established T-shaped pattern reported for Westerners observers, which is merely the result of the group averaging of the idiosyncratic visual sampling strategies of individual observers (Mehoudar et al., 2014).

Despite the growing literature on the existence of idiosyncratic sampling strategies, their functional role and the underlying neural mechanisms remain poorly understood. Some studies have investigated the impact of the fixated facial information input on neural responses, by recording the electroencephalographic (EEG) signals while observers fixated different facial information (i.e., viewing positions; VPs). This body of work has focused on the N170 ERP (Event Related Potential) component (Bentin, Allison, Puce, Perez, & McCarthy, 1996), the earliest face sensitive neural marker characterized by an occipito-temporal negative deflection of the EEG signal 170ms after stimulus onset. Collectively, the results of these studies demonstrated that VPs differentially modulate the N170, with fixation on the eye region electing larger amplitudes (de Lissa et al., 2014, Itier, Latinus, & Taylor, 2006; Nemrodov, Anderson, Preston, & Itier, 2014; Rousselet, Ince, van Rijsbergen, & Schyns, 2014). This observation would suggest a possible universal *neural* preference toward this facial information. However, these studies have mainly involved grand-average analyses and did not control for individual fixation preferences. As a consequence, while this analytical approach allows enhancing commonalities across observers, it confounds crucial individual differences, leaving unaddressed the question of whether idiosyncratic fixation biases concur with idiosyncratic neural responses.

Fast-periodic visual stimulation (FPVS) has been increasingly used to examine different aspects of face processing, including e.g. face detection, discrimination, and categorization (Ales, Farzin, Rossion, & Norcia, 2012; Norcia, Appelbaum, Ales, Cottereau, & Rossion, 2015; Rossion, Torfs, Jacques, & Liu-Shuang, 2015; Liu-Shuang, Norcia, & Rossion, 2014). Compared to traditional ERPs, the FPVS response has many advantages: it is far less susceptible to noise artefacts, and its remarkably high signal-to-noise ratio increases the likelihood of detecting subtle differences between experimental manipulations of interest (Norcia et al., 2015). Such signal properties make the FPVS paradigm paired with EEG recordings ideal to investigate the potential relationship between VP dependency of neural responses and idiosyncratic visual sampling strategies.

To this aim, we tracked the eye movements of observers performing an old/new face recognition task (Blais et al., 2008) and extracted their fixation patterns. Within the same testing session, we then recorded their neural face discrimination responses by means of a FPVS paradigm, while observers fixated faces on one out of ten viewing positions covering all inner facial features (Fig. 2A). We then applied a robust statistical data-driven approach to relate the idiosyncratic sampling strategies and the electrophysiological responses across all electrodes independently, without any a-priori assumptions regarding the topography of a potential effect. To account for visual sampling idiosyncrasies, this computation was performed at the individual level. Our data show a strong positive relationship between idiosyncratic sampling strategies and neural face discrimination responses as recorded at different viewing positions, for *all* the observers. In particular, independently of the sampling strategy, the longer a viewing position was fixated under natural viewing conditions, the more likely this VP was to elicit the strongest neural face discrimination response when its fixation was enforced.

**Fig. 2.**
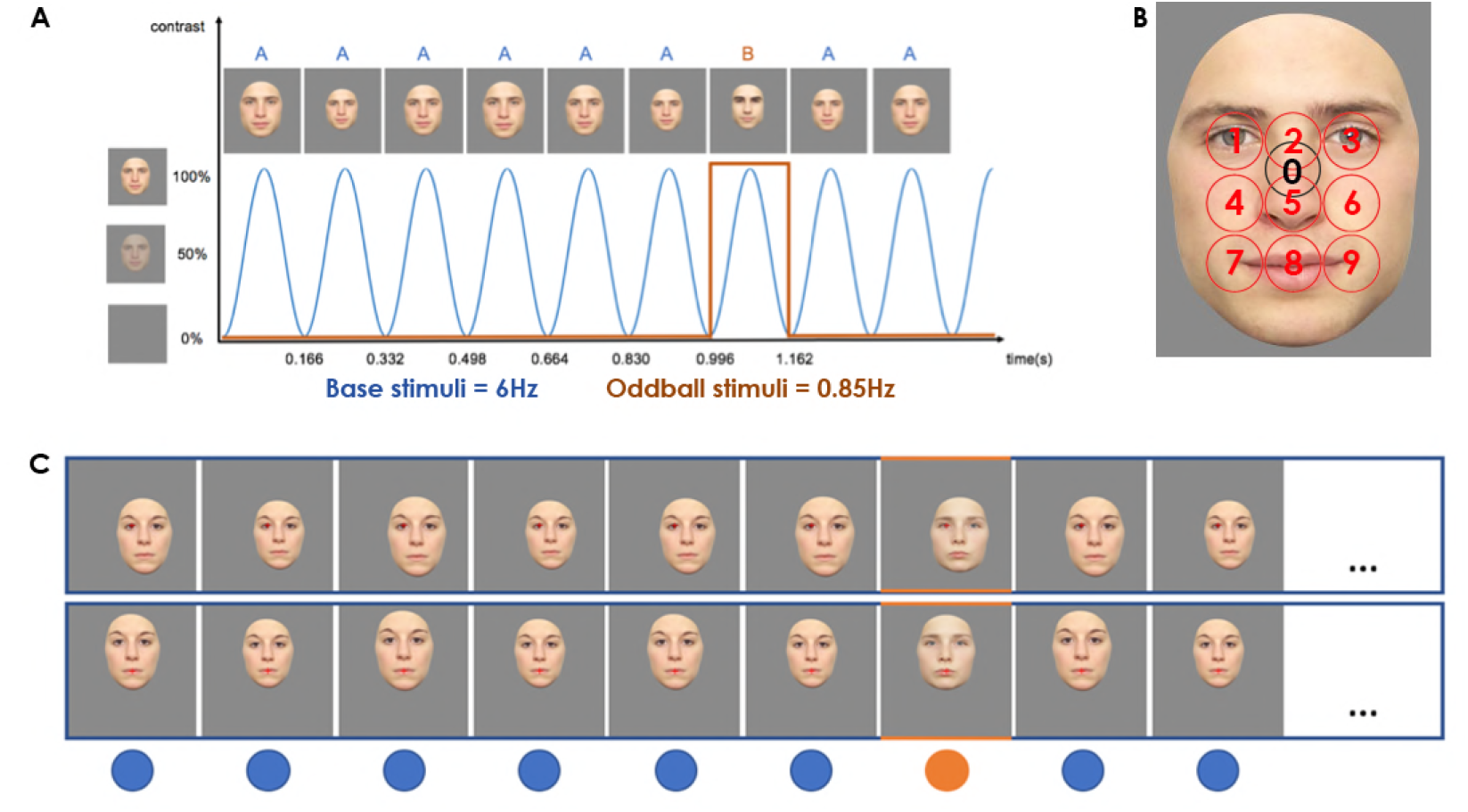
FPVS paradigm and viewing positions. (A) Faces were presented at a frequency rate of 6Hz through sinusoidal contrast modulation. Base stimuli consisted of images of the same facial identity; interleaved oddball stimuli conveying different identities were presented every 7^t^^h^ base stimulus. (B) Illustration of the 10 viewing positions (VPs) fixated by participants. (C) Examples of two trials displaying fixation on the left eye (VP1, top row), or mouth (VP8, bottom row).

## METHODS

### Participants

The sample size opted for was motivated by studies using the same FPVS paradigm to index neural face discrimination that were published up to data acquisition (Dzhelyova & Rossion, 2014a, 2014b; Liu-Shuang et al., 2014; Liu-Shuang, Torfs, & Rossion, 2016; sample size range: 8-12). In Dzhelyova and Rossion’s (2014b) study using a within subject design, the observed minimal effect size resulting from a repeated ANOVA was .2 (partial-eta). As the effect size estimation is often overly optimistic in the literature, we planned our experiment based on an effect size of .1 and an estimated sample size of 15 participant which results in a power of .95 to detect an effect. Based on prior experience and the requirement of high quality data from independent methods, we chose to test a total number of 20 participants. Our cohort comprised 20 Western Caucasian observers (11 females, two left-handed, mean age: 25±3 years) with normal or corrected-to-normal vision and no history of psychiatric or neurological disorders. Three observers were excluded due to poor quality of the eye movement data. All participants provided written informed consent and received financial compensation for participation; all procedures were approved by the local ethics committee.

### Procedures

### Eye-tracking

#### Stimuli and procedure

Stimuli consisted of 112 grey-scaled pictures portraying 56 Western Caucasians (i.e. WC) and 56 East Asians (i.e. EA) respectively obtained from the KDEF (Lundqvist, Flykt, & Öhman, 1998) and the AFID (Bang, Kim, & Choi, 2001). Faces were presented at a viewing distance of 75 cm and subtended 12.56° (height from chin to hairline) × 9.72° (width) of visual angle on a VIEWPIxx/3D monitor (1920 × 1080 pixel resolution, 120 Hz refresh rate).

Participants performed an Old-New face recognition task (Blais et al., 2008), with two blocks, each comprising a learning and a recognition phase of either WC and EA faces. In each learning phase participants were presented with 14 identities (7 female) with a neutral, happy or disgust expression; the recognition phase involved the presentation of these encoded identities alongside of 14 new ones, with a change of facial expression for the learned identities in order to prevent for face image matching strategies instead of genuine face recognition. Participants were required to indicate via button press whether a stimulus had been previously seen or not. During the learning phase, the faces were presented for five seconds; during the recognition phase presentation was terminated upon participants’ responses. The eye movements were recorded during both the learning and recognition phases.

#### Data acquisition and processing

The oculomotor behavior was recorded for each participant using an EyeLink 1000 Desktop Mount with a temporal resolution of 1000 Hz. The raw data are available in the public domain (Stacchi, Ramon, Lao, & Caldara, 2018). Data were registered by using the Psychophysics (Brainard, 1997) and the EyeLink (Cornelissen, Peters, & Palmer, 2002) Toolbox running in a MatlabR2013b environment. Calibrations and validations were performed at the beginning of the experiment using a nine-point fixation procedure. Additionally, before each trial a fixation cross appeared in the center of the screen and participants were instructed to fixate on it until a new stimulus appeared to ensure eye movements were correctly tracked. A new calibration was performed if the eye drift exceeded 1° of visual angle.

After removing eye blinks and saccades using the algorithm developed by Nystrom et al. (Nyström & Holmqvist, 2010), observers’ eye movement data from the learning phases of the Old-New task were processed to create individual fixation maps. Previous studies have shown that with this paradigm there are no differences in the sampling strategies used to samples WC or EA faces (Blais et al., 2008; Caldara, 2017). Therefore, in order to increase the signal-to-noise ratio, fixation maps were extracted from all face presentations. Individuals’ fixation intensities (based on the cumulative fixation duration) per observers were derived using these fixation maps and pre-defined circular regions of interest (ROIs; see Fig. 1). The ROIs covered 1.8° of visual angle and were centered on the ten viewing positions fixated during the FPVS experiment.

**Fig. 1.**
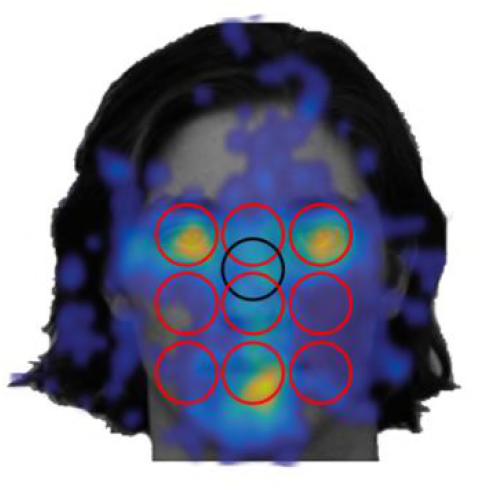
Illustration of the regions of interest (ROI) surrounding the 10 viewing positions. Observers’ fixation maps were overlaid onto a ROI mask to compute the fixation intensity per ROI. The ROI were covering 1.8° of visual angle and were centered on 9 equidistant viewing positions (red circles) and on an additional VP corresponding to the center of the stimulus (black circle).

### EEG

#### Stimuli and procedure

We used full-front, color images of 50 identities (25 female) from the same set described previously (Liu-Shuang et al., 2014). All faces conveyed a neutral expression, were cropped to exclude external facial features, and were presented against a grey background. Each original stimulus subtended 11.02° (height) × 8.81° (width) of visual angle at a viewing distance of 70 cm.

Face stimuli were presented at a constant frequency of 6 Hz, with intervening oddball identities every 7^th^ base (0.857 Hz) (Fig. 2A). The experiment comprised 20 trials: ten conditions (the viewing positions participants were required to fixate on; Fig. 2B), with two 62s trials per condition (trials differed with respect to the gender of the face stimuli). To prevent eye-movements, participants were instructed to maintain fixation on a central cross. The position of face stimuli was manipulated to vary, across trials, the fixated viewing position, hence the facial information. Faces were presented through sinusoidal contrast modulation (see Fig. 2A). Additionally, two seconds of gradual fade in and fade out were added at the beginning and end of each trial. To maintain subjects’ attention, the fixation cross briefly (200ms) changed color (red to blue) randomly between seven and eight times within each trial; participants were instructed to report the color change by button press. Subjects were also monitored trough a camera placed in front of them communicating the experimenter computer. Finally, to avoid pixel-wise overlap, stimulus size varied randomly between 80% and 120% of the original size (visual angle ranged between 8.82-13.22° (height) and 7.05-10.57° (width)).

#### Data acquisition and processing

Electrophysiological responses were recorded with Biosemi Active-Two amplifier system (Biosemi, Amsterdam, Netherlands) with 128 Ag/AgCl active electrodes and a sampling rate of 1024Hz. Additional electrodes placed at the outer canthi and below both eyes registered eye movements and blinks; electrode impedance was maintained between ±25kΩ. The recorded EEG was analyzed using Letswave 5 (http://nocions.webnode.com/letswave; (Mouraux & Iannetti, 2008)). The raw data are available in the public domain (Stacchi et al., 2018). Preprocessing consisted in high-and low-pass filtering the signal (with a 0.1Hz and 100Hz Butterworth band-pass filter (4^th^ order). Data were subsequently downsampled to 256Hz and segmented according to condition resulting in 20 66-second epochs, which included two seconds before and after stimulation. Independent component analysis was performed on each participant’s data to remove contamination due to eye-movements.

Noisy electrodes were interpolated using the three nearest spatially neighboring channels; this process was applied to no more than 5% of all scalp electrodes. Segments were then re-referenced to a common average reference and cropped to an integer number of oddball cycles, excluding two seconds after stimulus onset and two seconds before stimulus offset (∼58-second epochs; 14932 bins). Epochs were then averaged separately for each subject per condition.

#### Frequency domain

Fast Fourier Transform (FFT) was applied to the averaged segments and amplitude was extracted. The data were baseline corrected by subtracting from each frequency’s amplitude the average of its surrounding 20 bins excluding the two neighboring ones. Finally, for each subject and condition, the summed baseline-corrected amplitude of the oddball frequency and its significant harmonics provided the index of neural face discrimination. Following previous procedures (Dzhelyova, Jacques, & Rossion, 2016), harmonics were considered significant until the mean z-score across all conditions was no longer above 1.64 (*p*<.05). Based on this criterion we considered the first 11 harmonics excluding the 7^th^ harmonic, which is confounded with the base stimulation frequency rate.

#### Analysis

Using the iMAP4 toolbox (Lao, Miellet, Pernet, Sokhn, & Caldara, 2017) we computed a linear regression to explore the relationship between fixation bias (the z-scored fixation duration) and neural face discrimination (i.e., the FPVS response amplitude) within observer. To this aim we performed a linear mixed-effects model with random effect for intercept and *Fixation duration* grouped by subject. To avoid a-priori assumptions regarding topography of the effect, we regressed the two variable at all scalp electrodes independently.

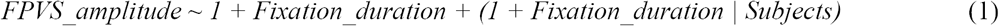

This computation will determine whether, VP-dependent fixation duration are associated with the amplitude of the neural face discrimination response elicited by each VP, for each subject independently. Importantly, because the analysis is performed at the individual level, there is no a priori expectation on how VPs are ranked. We opted for this approach in light of individual differences in fixation patterns reported previously (Mehoudar et al., 2014; Arizpe et al., 2016; Kanan et al., 2015), and similar idiosyncrasies assumed to exist for neural face discrimination responses across VPs. Therefore, the model used here allows each subject to have his/her specific VP-pattern and a relationship emerges if the fixation pattern is predictive of the neural response pattern of the same subject. Finally, as the current work does only focus on individual subjects, we did not perform any analysis involving average fixation maps and average EEG responses. The processed data used to compute this analysis are available in the public domain (Stacchi et al., 2018)

## RESULTS

### Behavior

As expected, subjects’ performance in the Old-New task, as indexed by d’, was significantly better for Western Caucasian (M=1.62, SD=.64) than East Asian faces (M=0.97, SD=.60), t(16)=5.72, *p*<.01. Subjects’ performance was nearly at ceiling for the FPVS orthogonal task consisting in detecting the color-change of the fixation-cross (M=.95, SD=.11). Due to technical issues, one subject’s behavioral response at this orthogonal task was not recorded. However, as for all subjects, behavior was monitored trough a webcam.

### Eye-movements and FPVS response

#### Description of fixation and neural biases at the group and individual level

The average fixation maps (computed for descriptive purposes and) shown in Figure 3A demonstrate that *as a group* observers preferentially sampled facial information encompassing the eyes, nasion, nose and mouth. However, because the focus of this work was to investigate the relationship between fixation patterns and neural responses at the individual level, group data were not subject to any further analysis.

**Fig. 3.**
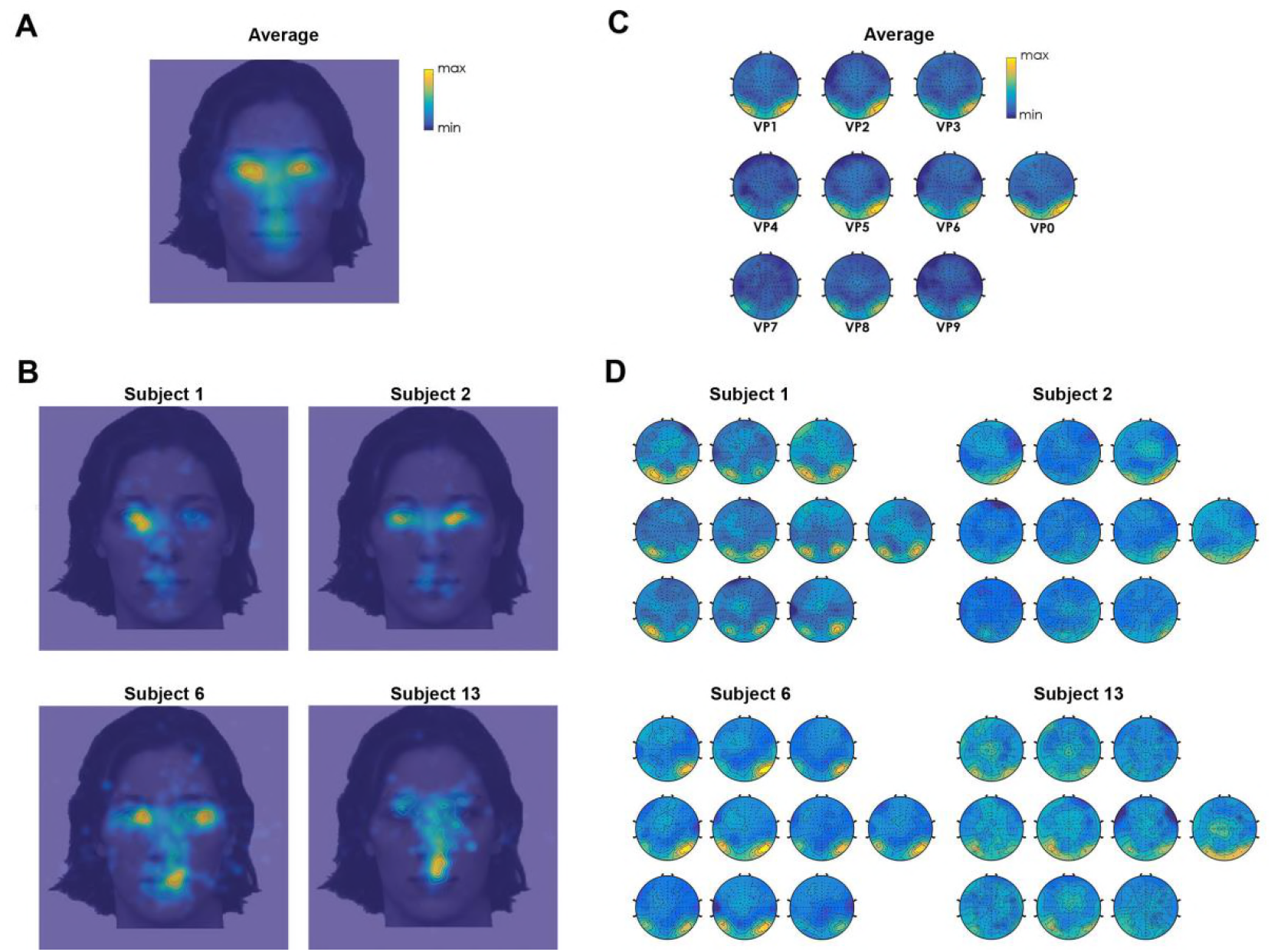
Fixation maps and oddball responses. A and C show the grand-average fixation map and FPVS responses respectively, while B and D show the two measures for the same subjects. For illustration, only four subjects are reported.

At the individual level, the majority of individual observers’ fixation maps did not *perfectly* conform to the grand average fixation pattern (Figure 3A-B – see also supplementary Figure S1), clearly demonstrating the existence of idiosyncratic visual sampling strategies (see Figure S1 for all individual observers’ fixation maps). Mirroring these results, the grand average neural face discrimination response amplitudes varied as a function of VPs, with the greatest amplitudes for the central position (Figure 3C). However, the neural response amplitudes also markedly differed across individuals (Figure 3D).

#### Regression analysis: Assessing the relationship between fixation and neural biases at the individual level

The data-driven regression between individuals’ fixation durations and FPVS responses across VPs computed independently on all electrodes revealed a positive relationship at right occipito-temporal (OT) and central-parietal (CP) clusters (see Figure 4A).

**Fig. 4.**
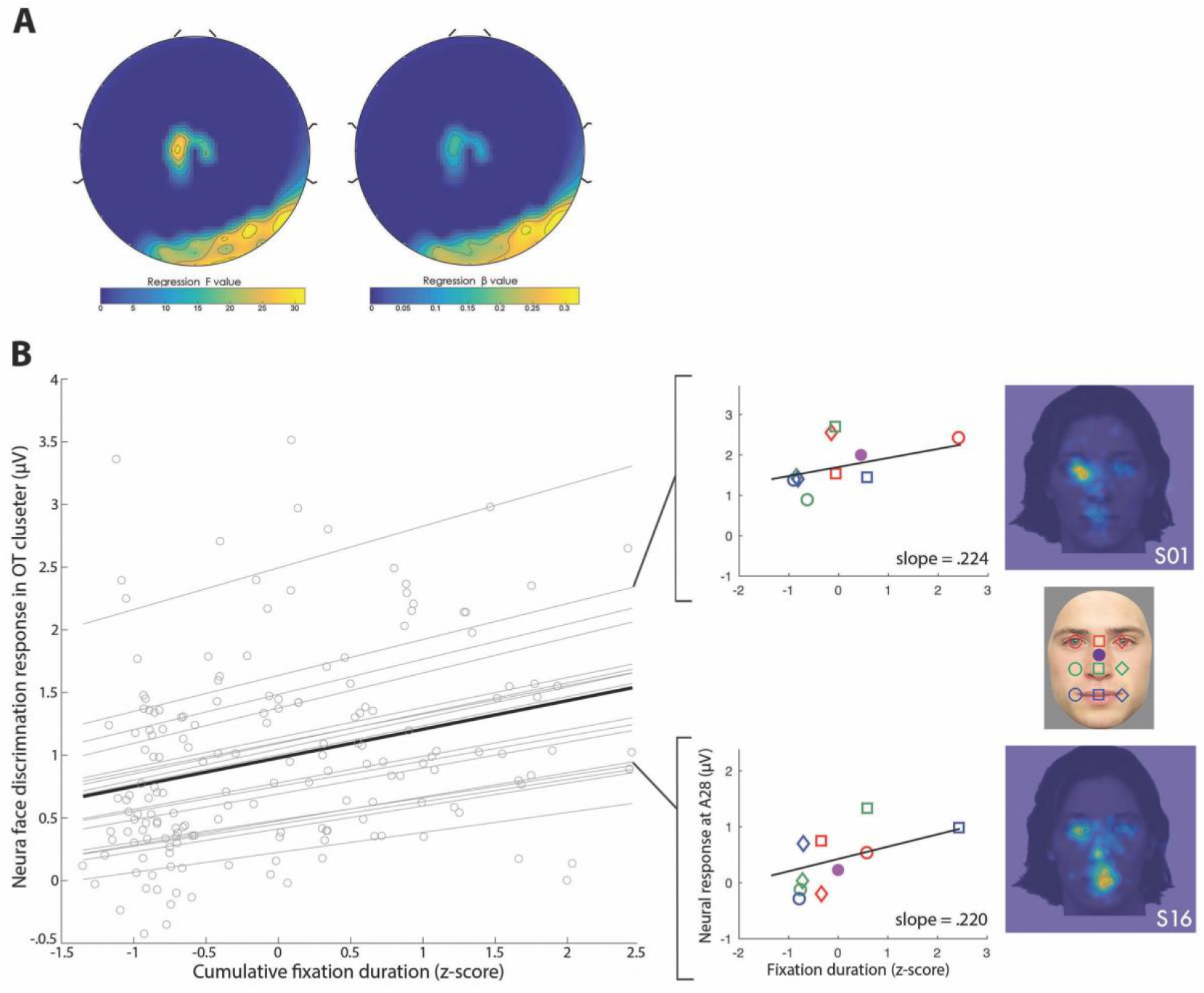
The relationship between fixation duration and neural face discrimination responses across VPs observed across all subjects considered individually. (A) Regression F-values (left) and beta maps (right). Original maps were overlaid with a mask and only electrodes exhibiting a significant effect (p< 7.81e-05) are shown. (B) On the left, the scatterplot illustrates individual subjects’ (light grey lines) as wells as the group (black line) effect averaged across the significant occipito-temporal cluster of electrodes. Crucially, the ranking of fixation and neural bias for VPs differed across subjects. On the right, data of two subjects illustrate that the relationship between fixation and neural bias emerges irrespectively of the fixation pattern exhibited (i.e., left-eye for S01 and mouth for S16). Note that here individual subjects’ correlations are displayed (black line).

The occipito-temporal cluster includes 12 significant electrodes with the strongest effect at A28 (F(1,169)=30.02, ß=.22 [.14 .30], *p*=1.54e-07) and the smallest at A13 (F(1,169)=17.18 ß=.20 [.10 .29], *p=*5.37e-05) (Table S1). The average across the electrodes in the OT significant cluster shows that individual subjects exhibited variable intercepts but similar slopes (see Fig. 4B). Despite inter-individual variations in the neural face discrimination response amplitude and fixation durations, the relationship was present within *all* the observers (see Fig. 5).

**Fig. 5.**
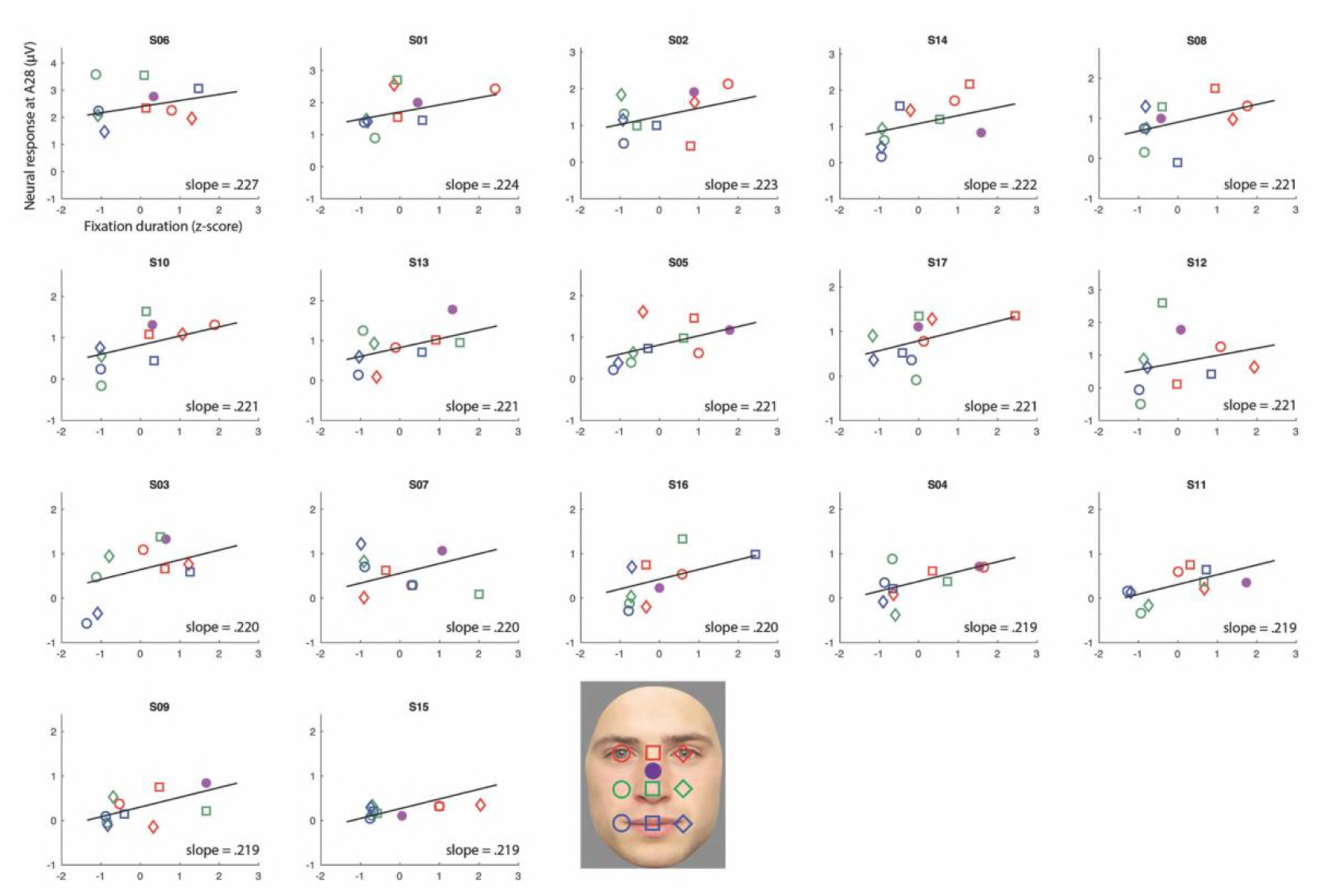
The relationship between individual subjects’ fixation and neural bias across VPs. For each observer (S1-S17), the VP-dependent fixation duration (x-axis) is plotted against their neural face discrimination response amplitude (y-axis) along with their individual correlation (black line); VPs are color- and shape-coded as indicated in the legend. The subjects are ordered as a function of their correlation strength, from high (.227) to low (.219). Although observers exhibited idiosyncratic VP-dependent fixation durations, all showed the general pattern, that facial features fixated longer (i.e., VPs) elicited larger neural responses. Note that here the neural face discrimination response magnitude is displayed at the electrode showing the largest effect (i.e., A28).

A small effect was also found on the central-parietal cluster comprising seven electrodes, with D14 showing the strongest effect (F(1,169)=26.12 ß=.13 [.08 .17], *p*=9.64e-07) and C1 exhibiting the smallest effect (F(1,169)=17.78 ß=.11 [.06 .17], *p*=4.05e-05) (Fig. 4A; Table S1).

#### Can **specific** fixation biases account for the observed relationship?

To assess whether subjects exhibiting a particular fixation bias (e.g., for the eyes) would show stronger correlations between fixation and neural biases, we first ranked observers’ fixation maps based on the magnitude of their individual relationship. As shown in Figure 6A, subjects showing similar fixation patterns could exhibit relationships of slightly different magnitude (e.g., left eye: S01 and S04), while observers exhibiting different fixation maps could show comparable correlation strengths (e.g., S10 and S13). Additionally, we computed the distance of each observer’s fixation map from the average fixation pattern. In this case, each map is treated as a vector and the measure of interest is the cosine distance between each observers’ map and the average one. This produces a value ranging between 0 and 1 for each subject. The higher the distance the more dissimilar that given subject’s pattern is from the average. Finally, we performed a Spearman correlation between this distance and the strength of the relationship between fixation and neural bias, which resulted to be non-significant (r=-.18, *p*=.49) (Figure 6B).

**Fig. 6.**
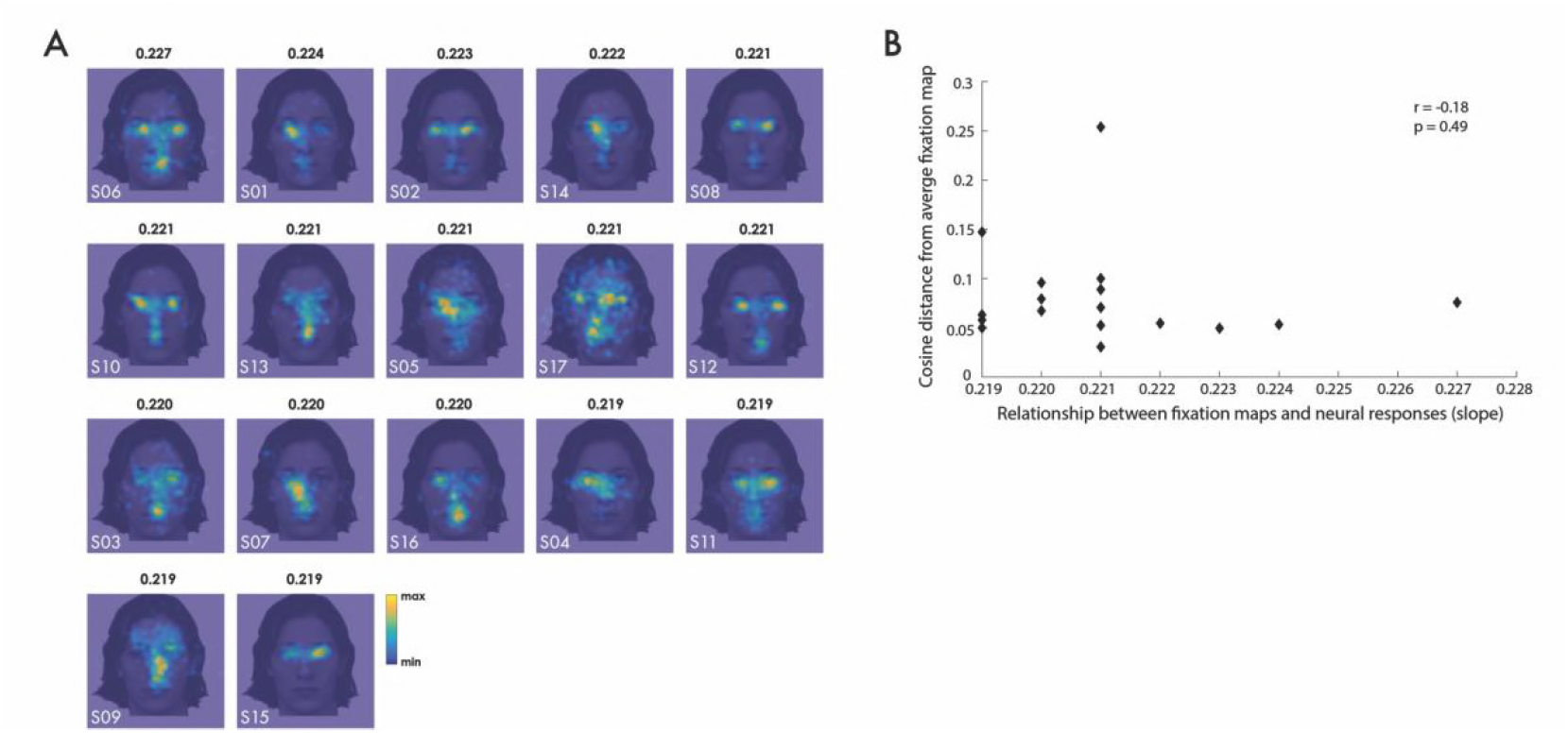
Fixation maps and strength of the fixation-neural-bias relationship. (A) Observers’ fixation maps sorted as a function of the slope of observers’ relationship between fixation bias and neural face discrimination response amplitude. The slope is reported for the electrode showing the strongest effect (i.e., A28). (B) The scatterplot illustrates the lack of correlation between: the cosine distance of individuals’ fixation maps from the average fixation map (y-axis) and strength of the relationship between fixation and neural bias (x-axis). Data show there was not a particular fixation bias more likely to correlate with the neural bias.

## DISCUSSION

This study investigated the relationship between *idiosyncratic* visual sampling strategies for faces and the magnitude of neural face discrimination responses during fixation on different types of facial information. Our data show that visual information sampling is distinct across observers, and these differences are positively correlated with *idiosyncratic* neural responses predominantly at occipito-temporal electrodes. Specifically, the Viewing Positions (VPs) that elicited stronger neural face discrimination responses coincided with the VPs that were more fixated under free-viewing conditions. Altogether, our data show that face processing involves idiosyncratic coupling of *distinct* information sampling strategies and *unique* neural responses to the preferentially sampled facial information.

For many years, the accepted notion in vision research was that face processing elicits a unique and universal cascade of perceptual and cognitive events to process facial identity, with particular importance ascribed to information conveyed by the eye region. For instance, eye movement studies have revealed a bias towards sampling of the eye region (Blais et al., 2008), the diagnosticity of which has been further documented by psychophysical approaches (e.g., Bubbles) (Gosselin & Schyns, 2001). Electrophysiological studies have also reported increased N170 magnitude during fixation on the eyes, compared to other information (de Lissa et al., 2014; Nemrodov et al., 2014). Collectively, these independent findings were taken to support the existence of a fixation and neural preference for the eye-region that is shared across *all* observers.

However, this idea has recently been challenged. For example, findings from eye movement studies emphasize idiosyncrasies in sampling preferences that are highly distinct from the group-average T-shaped pattern (Arizpe et al., 2017; Mehoudar et al., 2014), or by the existence of cultural differences (Blais et al., 2008; Caldara, 2017). These individual differences are not systematically associated with performance, as “mouth lookers” (i.e., observers showing preferential fixation on the mouth) could perform similarly to “eyes lookers”. Equally, two “eyes lookers” could exhibit very different performance (Peterson & Eckstein, 2013). Nonetheless, each observer’s adopted sampling strategy is optimal in the sense that performance is maximal when fixation is enforced on preferably sampled information, and decreases during fixation of other information (Peterson & Eckstein, 2013). These results suggest that individual differences do not reflect random inter-subject variation, but rather subtend functional idiosyncrasies in face processing.

Our results replicate and extend these previous findings, by showing that idiosyncratic visual sampling strategies strikingly mirror individuals’ patterns of neural face discrimination responses across VPs. Specifically, the facial regions preferentially sampled during natural viewing were those eliciting stronger neural face discrimination responses when fixated. This pattern was present in all observers, with even some of them (n=4) showing a perfect match between the most fixated facial feature and the one eliciting the strongest neural response at the electrode showing the strongest statistical relationship.

The strong and striking relationship between information sampling and neural idiosyncrasies suggests a functionally relevant process. Eye movements feed the neural face system with the diagnostic information in order to optimize information processing. The eyes constantly move to center elements of interest in the fovea, where visual acuity is greatest. This critical functional role, coupled with the relationship reported here between idiosyncratic sampling strategies and the neural face discrimination response pattern thus leads to two main considerations. First, our data show that face identity processing involves a fine-tuned interplay between oculomotor mechanisms and face-sensitive neural network. Second, the diagnosticity associated different types of facial information varies across observers. For a long time, researchers have debated on the nature of face representations, mainly opposing the idea of faces being represented as indivisible wholes (holistic or configural), as opposed to a collection of multiple, distinctively perceivable features (featural). This ongoing debate cannot be settled based on our finding of visual and neural idiosyncrasies. These idiosyncrasies do, however, refute the concept of a *single* face representation format shared across observers.

Our observations raise further important methodological and theoretical questions. The first concerns the traditional approach of standardizing the visual input to allow comparability across observers. This inherently creates a bias as not all observers are comparably tuned to this conventional VP. Additional open questions concern for instance (a) the extent to which the relationship between the visual sampling strategies and neural response patterns is *task*-specific, and (b) the direction of this relationship. Future studies are also necessary to establish whether similar effects can be found for scene, object and word processing. Finally, our approach may offer a promising novel route in clinical settings, if disorders comprising face processing impairments (i.e., prosopagnosia, autism, schizophrenia, etc.) involved an abnormal relationship between fixation patterns and neural responses to faces.

## CONCLUSION

When processing faces, observers deploy idiosyncratic sampling strategies by preferentially moving their eyes towards specific features. To assess the neural correlates underlying these idiosyncrasies, we recorded eye movements and neural face discrimination responses by means of FPVS. Our data show that idiosyncratic sampling strategies are mirrored by neural preferences, as facial information that is sampled longer during free viewing elicits stronger neural face discrimination responses when fixation is enforced on this feature. These findings show that eye movements and neural responses are finely tuned towards the facial information that is optimal for *individual observers.* The biologically relevant feat of processing facial identity is achieved by the use of idiosyncratic visual and neural representations.

## SUPPLEMENTAL INFORMATION

**Fig. S1.**
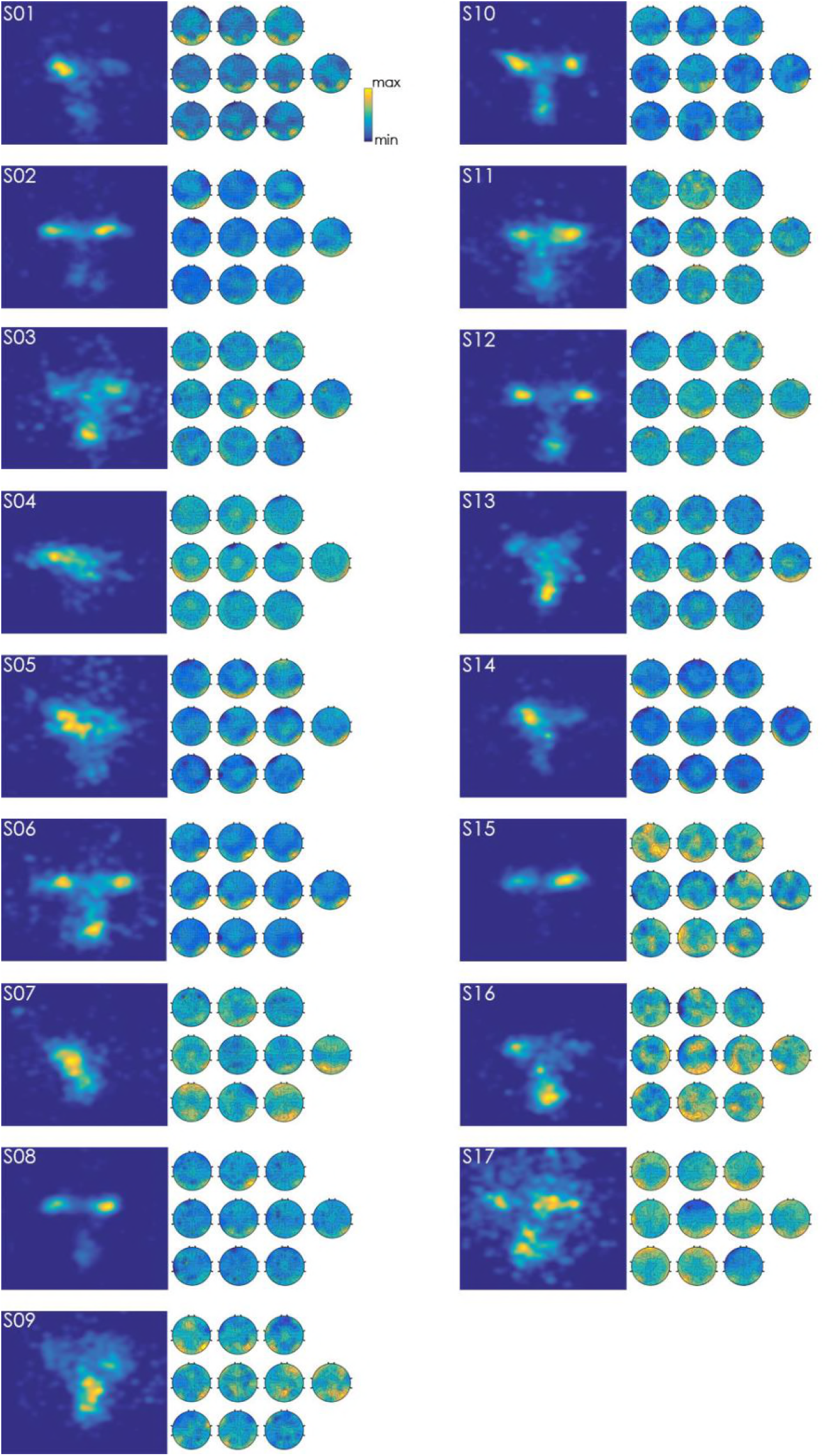
Fixation maps and neural face discrimination responses of all subjects.

**Table S1.**
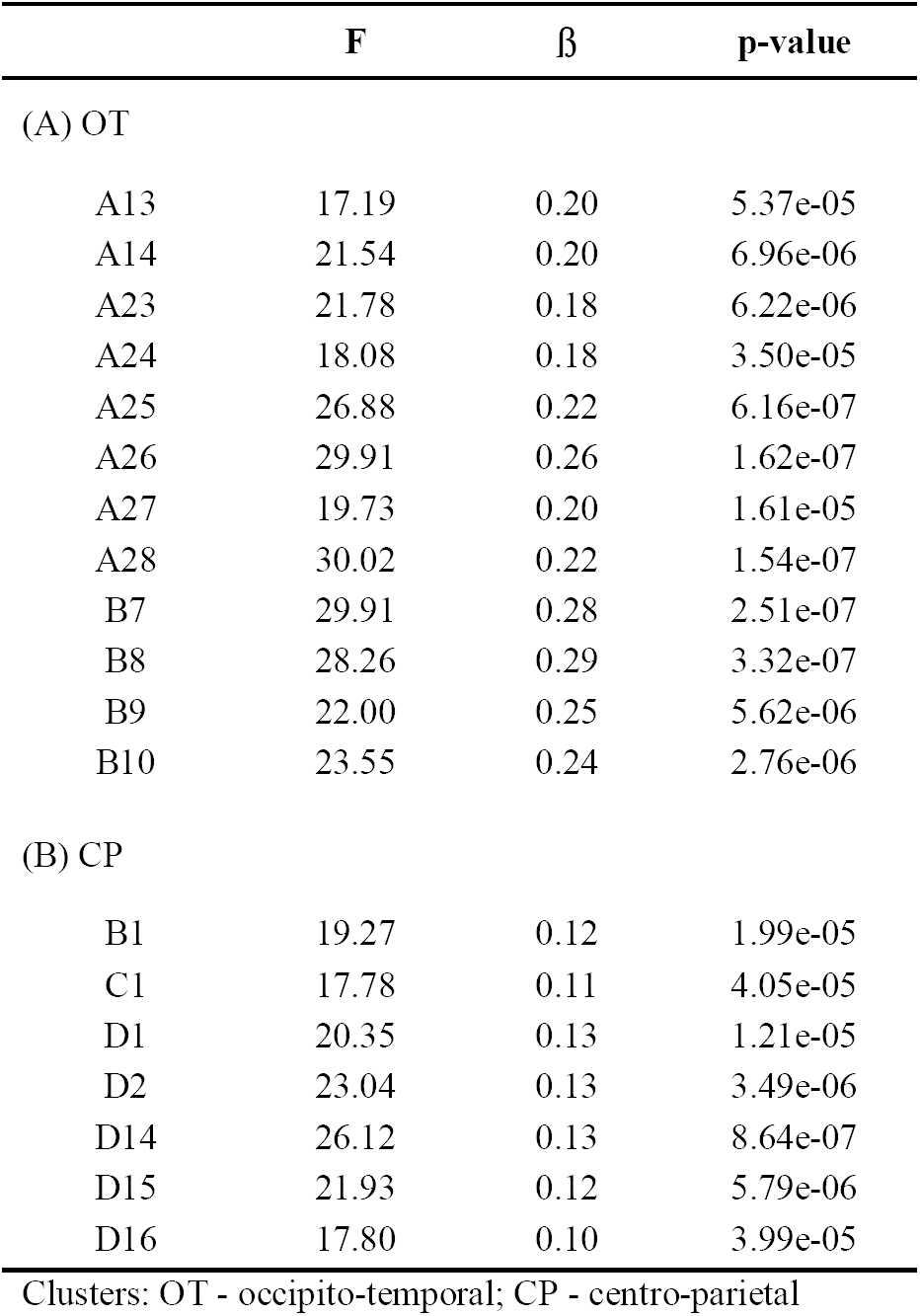
F, beta and p-values for electrodes showing a significant effect.

|  | **F** | **β** | **p-value** |
| --- | --- | --- | --- |
| **(A) OT** |  |  |  |
| A13 | 17.19 | 0.20 | 5.37e-05 |
| A14 | 21.54 | 0.20 | 6.96e-06 |
| A23 | 21.78 | 0.18 | 6.22e-06 |
| A24 | 18.08 | 0.18 | 3.50e-05 |
| A25 | 26.88 | 0.22 | 6.16e-07 |
| A26 | 29.91 | 0.26 | 1.62e-07 |
| A27 | 19.73 | 0.20 | 1.61e-05 |
| A28 | 30.02 | 0.22 | 1.54e-07 |
| B7 | 29.91 | 0.28 | 2.51e-07 |
| B8 | 28.26 | 0.29 | 3.32e-07 |
| B9 | 22.00 | 0.25 | 5.62e-06 |
| B10 | 23.55 | 0.24 | 2.76e-06 |
| **(B) CP** |  |  |  |
| B1 | 19.27 | 0.12 | 1.99e-05 |
| C1 | 17.78 | 0.11 | 4.05e-05 |
| D1 | 20.35 | 0.13 | 1.21e-05 |
| D2 | 23.04 | 0.13 | 3.49e-06 |
| D14 | 26.12 | 0.13 | 8.64e-07 |
| D15 | 21.93 | 0.12 | 5.79e-06 |
| D16 | 17.80 | 0.10 | 3.99e-05 |

Clusters: OT - occipito-temporal; CP - centro-parietal

